# Single-Strand DNA Breaks Cause Replisome Disassembly

**DOI:** 10.1101/2020.08.17.254235

**Authors:** Kyle B. Vrtis, James M. Dewar, Gheorghe Chistol, R. Alex Wu, Thomas G. W. Graham, Johannes C. Walter

## Abstract

DNA damage impedes replication fork progression and threatens genome stability. Upon encounter with most DNA adducts, the replicative CMG helicase (<u>C</u>DC45-<u>M</u>CM2-7-<u>G</u>INS) stalls or uncouples from the point of synthesis, yet CMG eventually resumes replication. However, little is known about the effect on replication of single-strand breaks or “nicks”, which are abundant in mammalian cells. Using *Xenopus* egg extracts, we reveal that CMG collision with a nick in the leading strand template generates a blunt-ended double-strand break (DSB). Moreover, CMG, which encircles the leading strand template, “runs off” the end of the DSB. In contrast, CMG collision with a lagging strand nick generates a broken end with a single-stranded overhang. In this setting, CMG translocates beyond the nick on double-stranded DNA and is then actively removed from chromatin by the p97 ATPase. Our results show that nicks are uniquely dangerous DNA lesions that invariably cause replisome disassembly, and they argue that CMG cannot be deposited on dsDNA while cells resolve replication stress.

**Highlights:**

- The structures of leading and lagging strand collapsed forks are different
- CMG passively “runs off” the broken DNA end during leading strand fork collapse
- CMG is unloaded from duplex DNA after lag collapse in a p97-dependent manner
- Nicks are uniquely toxic lesions that cause fork collapse and replisome disassembly

## Introduction

Single-stranded DNA breaks or “nicks” are generated by ionizing radiation, free radicals, topoisomerase I, and as intermediates in base excision repair (BER) (Caldecott, 2008). Nicks encountered in the leading strand template during eukaryotic replication cause replication fork “collapse” with formation of a single-ended double-strand break (seDSB; Figure S1A) (Nielsen et al., 2009; Strumberg et al., 2000), but whether this also occurs at lagging strand nicks has not been examined. seDSBs can be repaired by a subpathway of homologous recombination (HR) known as break-induced replication (BIR), which involves resection of the broken end, invasion into the sister chromatid, and replication to the end of the chromosome or until a converging fork is encountered (Haber, 1999; Mayle et al., 2015). Loss of the recombination proteins RAD51, BRCA1, or BRCA2 is lethal in unperturbed vertebrate cells (Hakem et al., 1996; Sharan et al., 1997; Sonoda et al., 1998; Tsuzuki et al., 1996), consistent with estimates that ~50 replication forks normally collapse in every S phase (Vilenchik and Knudson, 2003). Moreover, cancer therapeutics such as topoisomerase and poly (ADP-ribose) polymerase (PARP) inhibitors function by stabilizing nicks and promoting replication fork collapse (Hengel et al., 2017). Thus, the repair of collapsed forks is essential for cell viability and represents a prominent target in cancer therapy.

A central question is whether the replicative helicase CMG participates in replication restart after fork collapse. Early work suggested that CMG participates in BIR and that it remains on chromatin during fork collapse (Hashimoto et al., 2012; Lydeard et al., 2010). Moreover, a recent single molecule study suggested that when replication forks encounter DNA damage, CMG moves onto parental DNA beyond the damage, and when repair is complete, it re-engages with the fork for replication restart (Wasserman et al., 2019). However, other studies conclude that CMG is not involved in BIR (Dilley et al., 2016; Natsume et al., 2017; Sonneville et al., 2019; Wilson et al., 2013). To determine whether CMG could function in replication restart, tracking its fate during replication stress in a physiological setting is crucial. Another key factor dictating repair is the DNA structure generated during fork collapse. The specific structure formed may depend on whether collapse occurs at a nick in the leading strand template (Figure S1A,i, hereafter “lead collapse”) or lagging strand template (Figure S1A,ii, hereafter “lag collapse”). Work in human cells suggests that during lead collapse, a blunt, seDSB is generated (Strumberg et al., 2000) (Figure S1A,i) that would have to undergo resection prior to strand invasion. However, the lag collapse fork structure is unknown. In summary, whether the replisome is recycled for BIR and which DNA structures and processing steps underlie this clinically relevant DNA repair pathway remain unclear.

Here, we use ensemble and single molecule approaches in *Xenopus* egg extracts to examine the fate of the replication fork after collision with strand-specific nicks. Strikingly, after both lead and lag collapse, the CMG helicase is lost from the DNA, but via distinct mechanisms. Further, analysis of the fork DNA structures generated shows that lag collapse is more amenable to repair than lead collapse. Our results establish replication fork collision with nicks as uniquely deleterious for replication fork progression but also indicate that there are important differences between lead and lag collapse.

## Results

### Lead collapse generates a nearly blunt seDSB and a gap in the lagging strand

We wanted to study DNA replication fork collapse at a site-specific nick using *Xenopus* egg extracts, which support sequence non-specific replication initiation on added DNA, followed by a complete round of replication (Figure S1B) (Walter et al., 1998). However, nicked plasmid added to egg extract was rapidly ligated (Figure S1C, lanes 1-6) before forks would be able to reach the nick. Chemical modification of nucleotides flanking the nick did not prevent ligation (data not shown). We therefore flanked the nick with Tet operator (*tetO*) sites to which we bound the Tet repressor (TetR) before adding the plasmid to extract. We reasoned that TetR might stabilize the nick by blocking access to DNA ligase. Indeed, although TetR did not block fork progression (see Figure 3D legend below), it increased the half-life of the nick ~45-fold (Figure S1C). To further increase the probability that forks encounter a nick, we used three consecutive nicks, each flanked by *tetO* sites (Figure 1A, inset). To compare fork collapse when nicks reside in the leading versus lagging strand templates, it was necessary to ensure that forks arrive at the nicks from only one direction. Therefore, we flanked the nicks on the right with an array of 48 Lac repressors (LacR) (Figure 1A), which blocks replication fork progression in egg extracts (Dewar et al., 2015; Duxin et al., 2014), preventing arrival of a second fork at the nick. In this configuration, lead collapse should occur when the rightward fork collides with a nick in the bottom strand (Figure 1A, inset).

**Figure 1.**
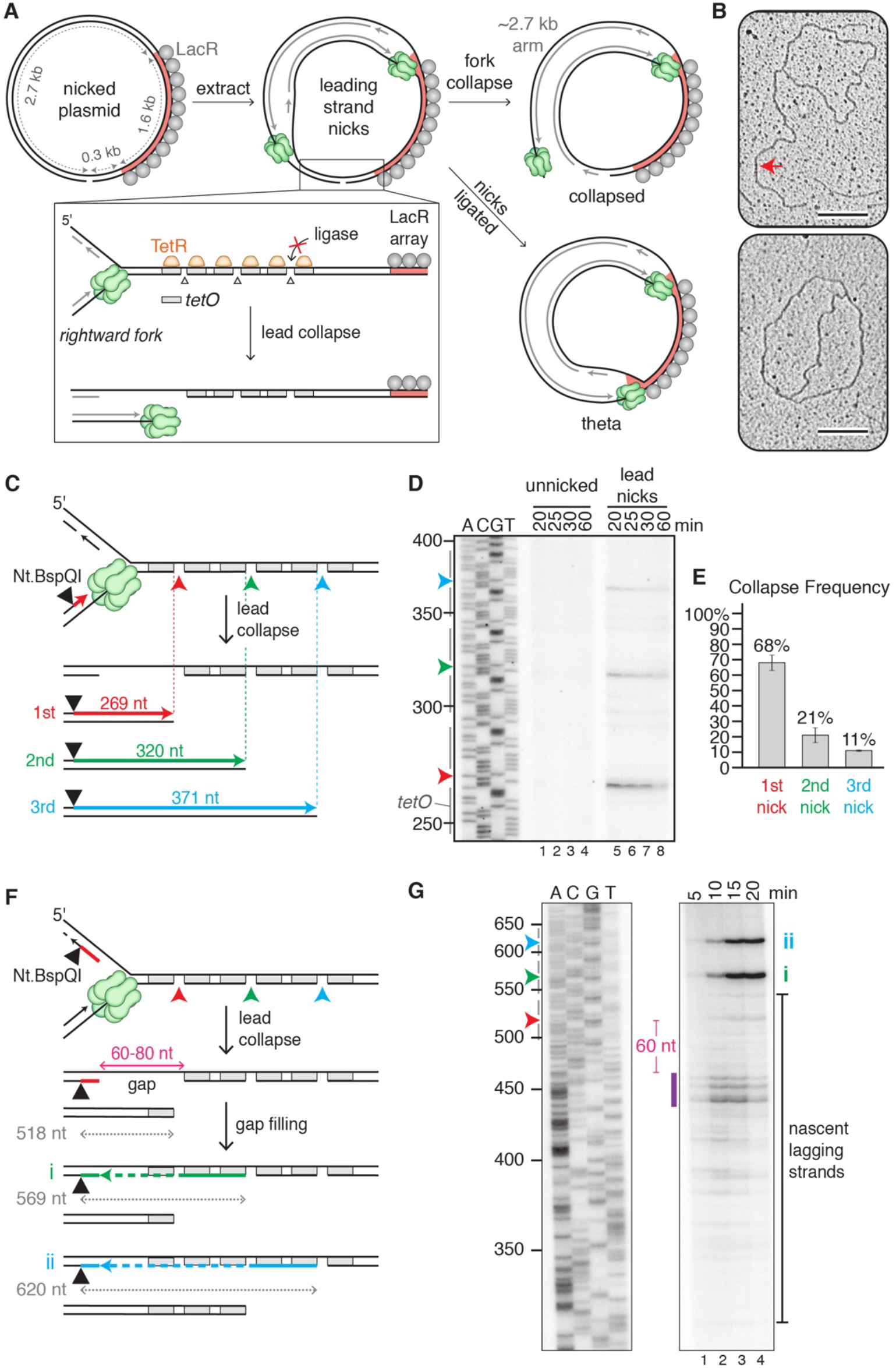
Lead collapse generates a nearly blunt seDSB and a gap in the lagging strand. (A) Depiction of experimental approach. (B) TEM images of collapsed (top) and theta (bottom) structures. Red arrow, 2.7 kb arm. Scale bar, 200 nm. (C and D) Depiction (C) and analysis (D) of nascent leading strands generated after lead collapse and DNA digestion with Nt.BspQI. Gray bars in (D) denote *tetO* sites. The DNA was visualized by autoradiography, and only the relevant segment of the gel is shown. (E) Percentage collapse at each of the three nicks out of total collapse events. Error bars, standard deviations from three independent experiments. (F) Repeat of (D), with Nt.BspQI site located on nascent lagging strand. After fork collapse at the first nick, gap filling from the 3’ end of the nick to the final Okazaki fragment and ligation should create a 569 nt product (i, green line). Collapse at the second nick, gap filling, and ligation should generate a 620 nt product (ii, blue line). (G) Nascent lagging strand analysis of products generated after lead collapse. Purple bar, most prominent lagging strand products. Irrelevant lanes between the two sets of lanes shown were removed. See also Figure S1.

When we replicated unnicked plasmid in extracts containing LacR and TetR, a prominent “theta” intermediate was generated (Figure S1D, lanes 1-3), as expected from forks stalling at the outer edges of the LacR array (Duxin et al., 2014). However, in the presence of a nick, a prominent new product appeared that migrated faster than theta (Figure S1D, lanes 4-6, red arrowhead). To identify the structure of this new product, we cut out the band, extracted the DNA, and performed transmission electron microscopy. This analysis revealed that the new product corresponds to a sigma-shaped, “collapsed” structure in which a linear arm of the expected size (2.7 kb; Figures 1A and S1E) is attached to the circular plasmid (Figure 1B, red arrow). Quantification showed that ~60% of the forks from the nicked plasmid underwent collapse (Figure S1D, lane 4). We attribute the remaining theta products (Figures 1A and 1B, bottom image; Figure S1D) to plasmids in which all three nicks were ligated before fork arrival. Our data provide direct visual confirmation of previous reports (Nielsen et al., 2009; Strumberg et al., 2000) that replication through a leading strand nick causes fork collapse with formation of a seDSB.

To determine the DNA structure of the broken end generated during lead collapse, we labeled nascent DNA stands with [α-^32^P]dATP and used the single strand endonuclease Nt.BspQI to cleave these strands at a defined position relative to the three TetR-flanked nicks (Figure 1C). Denaturing gel analysis showed that replication through the nick generated three ssDNA products whose 3’ ends were located close to each of the three nicks (Figure 1D, lanes 5-8). The product associated with collapse at the first nick was most abundant (Figure 1E), as expected from rightward fork movement. Higher resolution mapping showed that leading strand synthesis mostly stopped 3 nt from the end of the break, which should leave a 3 nt, 5’ single-stranded DNA overhang (Figures S1F and S1G, lanes 5-8). This interpretation is consistent with the properties of purified pol ε, the leading strand polymerase (Hogg et al., 2014). Our results suggest that lead collapse repair should generally require resection of the seDSB.

To map the distribution of nascent lagging strands during lead collapse, we next placed the Nt.BspQI site on the other strand (Figure 1F). Most nascent lagging strands were 440-460 nts in length (Figure 1G; purple bar). Because the first nick is located 518 basepairs from the Nt.BspQI site, we conclude that there is typically a ~60-80 nt gap between 5’ end of the nascent lagging strand and the site of collapse (Figure 1F, “gap”). By 20 minutes, lagging strand fragments declined, indicating gap filling (Figure 1G, lanes 3-4). Consistent with this interpretation, well-defined bands appeared that correspond to gap-filled products that retain the second and third nicks due to TetR (Figures 1F and 1G,i,ii). In summary, lead collapse generates a seDSB with a three nucleotide 5’ overhang and a lagging strand gap that is subsequently filled in. The data suggest that lead collapse repair involves not only resection but also gap filling.

### CMG is lost at the nick after lead collapse

Given that the leading strand is extended to within 3 nucleotides of the break, and because the CMG footprint on DNA comprises 20-40 nt (Fu et al., 2011), our data suggest that CMG loses its association with the DNA end during lead collapse. However, whether CMG dissociates altogether was unclear. To address this question, we used single molecule imaging to visualize CMG dynamics during lead collapse. We replicated stretched, immobilized DNA in GINS-depleted extract containing Alexa Fluor 647-labeled recombinant GINS (GINS^AF647^), a subunit of CMG (Figure S2A). Extracts also contained fluorescent Fen1 (Fen1^mKikGR^) to image nascent DNA synthesis (Loveland et al., 2012). As reported previously (Sparks et al., 2019), total internal reflection fluorescence microscopy revealed GINS^AF647^ molecules moving at the leading edge of growing Fen1^mKikGR^ tracts, demonstrating that GINS^AF647^ travels with active replication forks (Figures 2A,i, Supplemental Video 1). To generate a fluorescently labeled nick, we used a point mutant of Cas9, H840A (”nCas9”), which selectively nicks the non-target DNA strand (Jinek et al., 2012). nCas9 promoted efficient replication fork collapse (compare Figure S2B to Figure S1D). Furthermore, nCas9 RNPs labeled at the 5’ end of the tracrRNA with Atto550 (nCas9^Atto550^; Figure 2B) bound specifically to the target site on the stretched DNA (Figure S2C). However, nCas9^Atto550^ dissociated at a significant rate upon exposure to extract (Figures S3D and S3E; (Wang et al., 2020)). Therefore, to increase the likelihood of fork collapse, nCas9^Atto550^ was targeted to 4 sites in the same strand located ~1 kb apart. In most cases, only 1-2 nCas9 molecules remained bound at the time of fork arrival. The cluster of nicks was positioned 5-8 kb from one end of the 30 kb DNA such that a fork traveling along the short arm underwent lead collapse when it collided with nCas9 (Figure 2A,ii, Supplemental Video 2).

**Figure 2.**
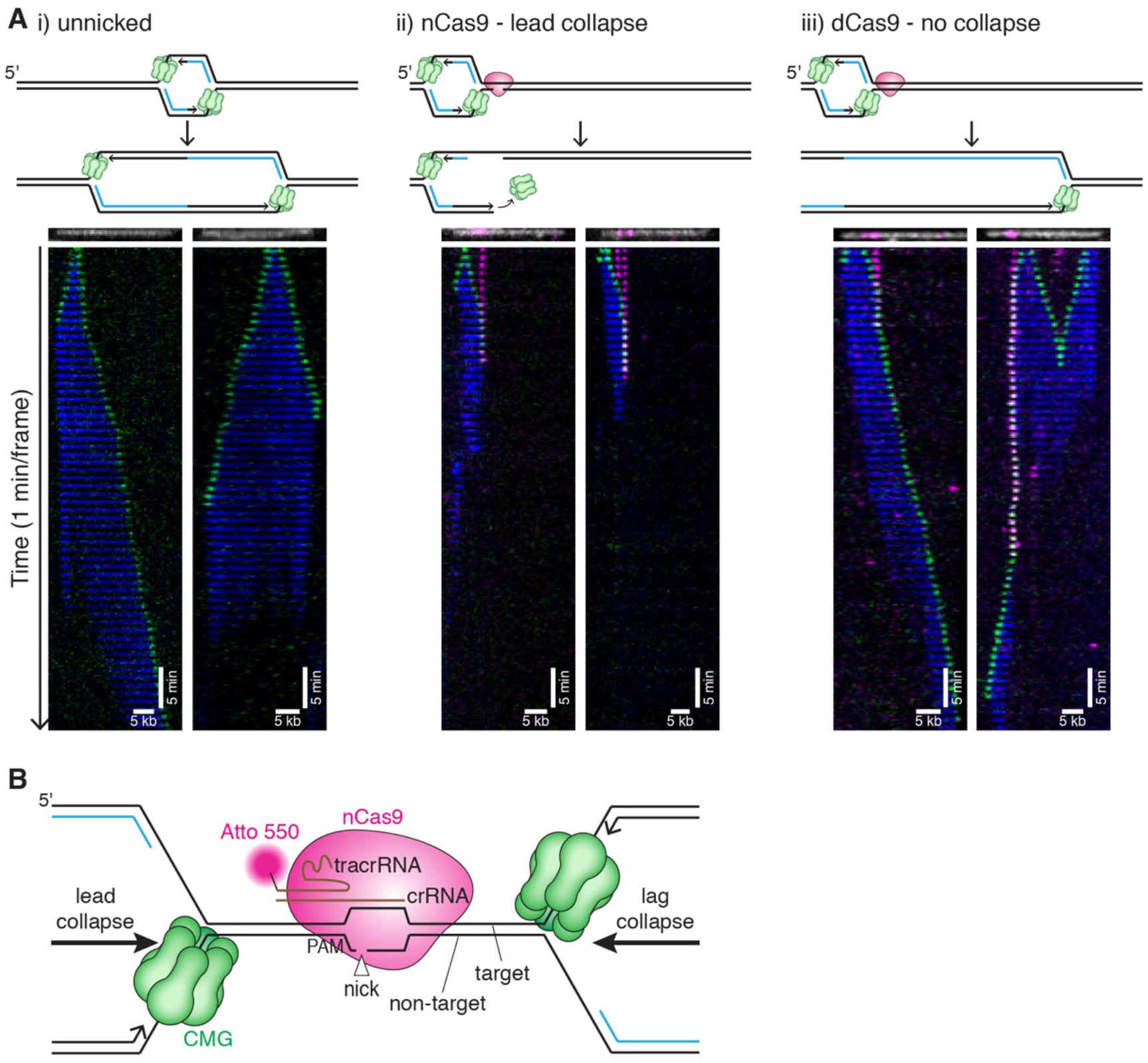
CMG is lost at the nick after lead collapse. (A) Representative single molecule kymographs and interpretive cartoons of i) replication on unnicked DNA, ii) lead collapse on nCas9-nicked DNA (note that in the kymograph on the right, the left nCas9 appears to dissociate before CMG arrival), and iii) replication fork encounter with dCas9. CMG is shown in green, Fen1^mKikGR^ in blue, nCas9 in magenta. Although more than one Cas9 is frequently bound, only one is depicted in cartoons for simplicity. Kymographs were created by stacking the frames of a movie (one-minute intervals). The DNA (white) and nCas9 (magenta) shown above each kymograph are from imaging the nCas9 and DNA before extract addition. See also Supplemental Videos 1-3. (B) Schematic of nCas9 H840A bound to DNA with replication forks arriving from the left (lead collapse) or right (lag collapse).

Representative examples of lead collapse are shown in Figure 2A,ii. In this scenario, CMG collided with nCas9, paused for a few minutes, and then disappeared. In 85% of such collisions, GINS dissociated from the chromatin at the nick site (Figure S2Fi,ii,vi). In contrast, when a catalytically dead mutant of Cas9 (dCas9) was targeted to these sites, CMG typically paused at dCas9 and then resumed replication (Figure 2A,iii, Figure S2G,iii, and Supplemental Video 3), indicating that CMG loss upon encounter with nCas9 is due to the nick. Consistent with a model in which CMG “runs off” the DNA end, GINS^AF647^ signal was lost at the same time as nCas9^Atto550^ 45% of the time (Figure S2F,i). Surprisingly, GINS^AF647^ was lost *before* nCas9^Atto550^ 33% of the time, suggesting that nCas9 may sometimes retain its interaction with DNA even after CMG slides off the end of the break (Figure S2F,ii). Such an outcome is plausible because CMG should be able to slide off the leading strand template without disrupting most of the contacts that nCas9 makes with the lagging strand template (Figure 2B, lead collapse). Instances in which CMG passes the nick site (Figure S2F,v) can mostly be attributed to the absence of a nick, as seen for dCas9, likely because nCas9 had not cleaved the DNA at the time of collision (see below). In summary, the loss of GINS^AF647^ at the nick site concurrent with or before nCas9 dissociation supports the model that CMG fully dissociates from the end of the break during lead collapse.

### Lag collapse generates a seDSB with a 3’ ssDNA overhang

What happens to the replication fork during lag collapse is unknown. To address this question, we replicated a plasmid with TetR-protected nicks on the lagging strand template (top strand) (Figure S3A). As shown in Figure S3B, this yielded the same new band observed during lead collapse (lanes 7-12). To determine the structure of the seDSB generated during lag collapse, we cleaved nascent lagging strands 515 bp from the first nick (Figure 3A). Denaturing gel electrophoresis showed that their 5’ ends were typically located 60-80 nucleotides from the break point (Figure 3B, purple bar), consistent with random initiation of the last Okazaki fragment relative to the nick. Therefore, lag collapse results in a seDSB break with a ~70 nt 3’ ssDNA overhang. Analysis of the nascent leading strands revealed that they were extended to within a few nucleotides of each nick (Figures 3C and 3D), and some were extended a few nucleotides beyond the nick, consistent with limited strand displacement synthesis (Figures S3C and S3D). Given CMG’s 20-40 nt footprint (Fu et al., 2011), this result argues that CMG does not stall when it hits the nick (Figure 3C,i) because no stalling products were detected 20-40 nt upstream of the nick (Figure 3D). Moreover, leading strand arrest at the nick also rules out that CMG continues unwinding DNA well beyond the nick (Figure 3C,ii). Instead, the data suggest that CMG either rapidly dissociates at the nick (Figure 3C,iii), or that it somehow translocates beyond the nick without unwinding DNA (Figure 3C,iv), either of which would allow leading strand synthesis to reach the nick.

**Figure 3.**
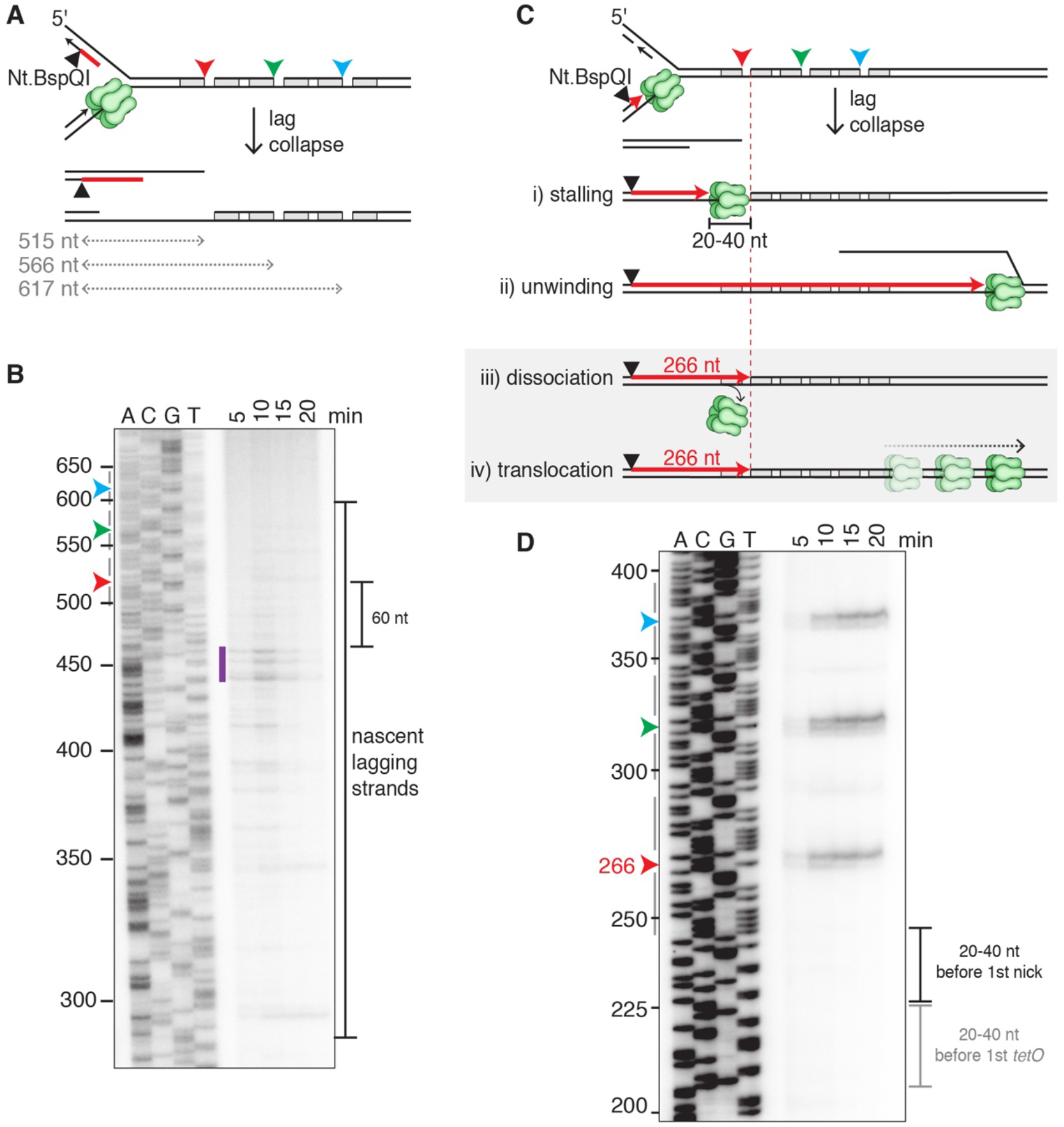
Lag collapse generates a seDSB with a 3’ ssDNA overhang. (A) Depiction of nascent lagging strand products generated from lag collapse at the first nick. 515 nt, 566 nt, and 617 nt correspond to the distances between the Nt.BspQI site and the three nicks. (B) Analysis of lagging strand products after lag collapse and DNA digestion with Nt.BspQI. Arrowheads, location of nicks. (C) Four different CMG fates after lagging strand fork collapse, including predicted size of the leading strand. Text for details. (D) Analysis of leading strand products after lag collapse and DNA digestion with Nt.BspQI. Black bracket, location where leading strands would stall in model (C, i). The lack of signal 20-40 nt before the 1^st^ *tetO* site (gray bracket) indicates that CMG does not stall at TetR.

### Active CMG unloading from double-stranded DNA during lag collapse

To distinguish between the latter two models, we examined the fate of fluorescent CMGs during lag collapse, which occurs when CMGs collide with nCas9^Atto550^ from the long arm of immobilized DNAs (Figure 4A, Supplemental Video 4). As seen for lead collapse (Figure 2A,ii), CMG typically paused at the nCas9, followed by rapid loss of nCas9 and CMG (Figure 4A, Veh.). However, far fewer CMGs were lost at the same time as nCas9 during lag collapse vs. lead collapse (4% vs. 45%; Figures. S2H,i and S2F,i), and more CMGs persisted longer than nCas9 for lag collapse compared to lead collapse (59% vs. 21%; Figures S2H,iii and S2F,iii). Together, our results demonstrate that CMG is lost from chromatin during lead and lag collapse, but the delayed CMG unloading during lag collapse could suggest a mechanism involving multiple steps.

**Figure 4.**
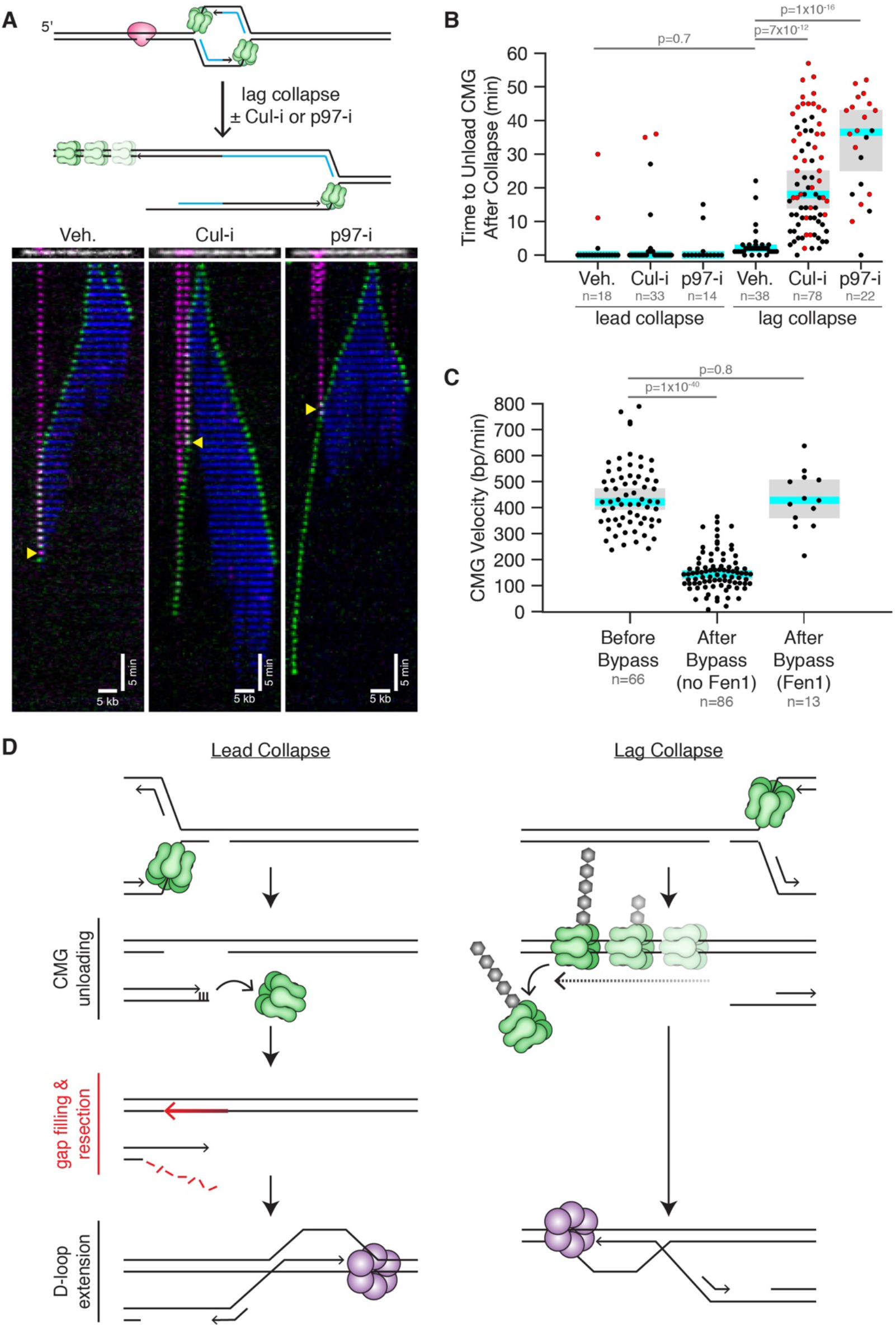
Active CMG unloading from double-stranded DNA during lag collapse. (A) Representative single molecule kymographs and interpretive cartoons when CMG approaches nCas9 from the long arm in the indicated conditions. Yellow triangle, moment of collapse (nCas9 loss). The DNA (white) and nCas9 (magenta) shown above each kymograph are from imaging the nCas9 and DNA before extract addition. See also Supplemental Videos 4-6. Note that in the middle kymograph, the left nCas9 appears to dissociate before arrival of CMG. (B) Distribution of CMG unloading times after lead and lag collapse in the indicated conditions. Because many CMGs persisted until the end of the 60 min experiment (red circles), the median times reported are lower limits. In (B) and (C), blue line is the median, and gray box represents the 95% confidence for the median determined from bootstrapping analysis. (C) Distribution of CMG velocities before and after nCas9 bypass with and without trailing Fen1 signal. (D) Models for lead and lag collapse. Purple hexamer, putative helicase that replaces CMG for BIR.

We hypothesized that CMG unloading during lag collapse might involve the same mechanism used during replication termination, in which converging CMGs are ubiquitylated by CRL2^Lrr1^ and unloaded by the p97 ATPase (Figure S4A) (Dewar and Walter, 2017). To test this idea, we supplemented extracts with MLN4924 (“Cul-i”), which inhibits all cullin-RING E3 ubiquitin ligases including CRL2^Lrr1^ (Dewar et al., 2017), or NMS873 (“p97-i”), an allosteric p97 inhibitor. Strikingly, Cul-i delayed CMG dissociation 9-fold after collision with lagging strand nicks, and the retained CMG continued unidirectionally translocating along the DNA away from the collapse site (Figures 4A and 4B, Supplemental Video 5). Similarly, p97-i inhibited CMG unloading and promoted continued translocation beyond the nick (Figures 4A and 4B, Supplemental Video 6). Importantly, Cul-i and p97-i did not affect CMG loss during lead collapse (Figure 4B). These results show that CMG unloading during lag collapse, but not lead collapse, involves a cullin-RING ligase and the p97 ATPase, as seen during termination.

CMG translocates onto dsDNA before being unloaded during termination (Dewar et al., 2015)(Low et al., in preparation). Our data suggest that a similar phenomenon occurs during lag collapse. When CMG translocated beyond a lagging strand nick in the presence of Cul-i or p97-i, no Fen1 signal was detected behind the helicase 85% and 84% of the time, respectively (Figure S2H,ix). Similarly, no Fen1 signal was observed when CMG translocated beyond the nick in the absence of Cul-i and p97-i (Figures S4C and S2H,ix), but such events were rare because CMG was usually unloaded before it could travel far enough to assess Fen1’s behavior (Figures S4D and S2H,viii). Thus, CMG translocation beyond the nick after lag collapse is not associated with DNA synthesis. Moreover, after colliding with a lagging strand nick, CMG translocated three-fold slower than before the collision (Figure 4C; 144 bp/min vs. 422 bp/min). However, the rate of CMG progression was the same before and after encounter with nCas9 in the few instances in which Fen1 signal did trail behind CMG after the encounter (Figures 4C and S4E). Thus, we infer that the fork failed to undergo collapse in these cases (Figures S2F,ix and S2H,x). In summary, after CMG travels beyond a lagging strand nick, it translocates slowly and does not promote DNA synthesis. This strongly implies that CMG transitions onto dsDNA at the nick, which is consistent with biochemical experiments using purified CMG (Kang et al., 2012; Langston and O’Donnell, 2017). Together, the data argue that during lag collapse, CMG is removed from dsDNA via the same mechanism that operates during replication termination (Figure S4A and S4B).

## Discussion

In this study, we have monitored the fate of the replisome at nicks, an abundant form of DNA damage. We find that when forks encounter nicks in the leading strand template, CMG undergoes passive dissociation when it slides off the end of the break; at lagging strand nicks, CMG transitions onto dsDNA and is recognized as a terminated complex, leading to active, p97-dependent removal. CMG anchors most proteins to the fork, suggesting that the entire replisome is effectively disassembled at both kinds of nicks. Importantly, there is no known mechanism to re-assemble CMG *de novo* in S phase. Therefore, our results strongly suggest that after fork collapse, resumption of DNA synthesis requires nucleation of a BIR replisome around a new DNA helicase. Our results are consistent with studies that BIR involves Pif1 in yeast and possibly MCM8-9 in mammalian cells (Natsume et al., 2017; Wilson et al., 2013). In summary, our data show that compared to most chemical adducts, strand discontinuities are uniquely dangerous DNA lesions that cause fork breakage and replisome disassembly.

The question arises whether CMG can be recycled during other forms of replication stress. Based on single molecule imaging with yeast CMG, it was proposed that when replication stalls at DNA damage, CMG diffuses onto dsDNA downstream of the damage; when repair is complete, CMG re-engages with the fork for replication restart (Wasserman et al., 2019). However, our data show that CMG is rapidly unloaded from dsDNA. Moreover, even if unloading could be prevented, CMG would translocate unidirectionally away from the stressed fork. Thus, our data suggest that in a cellular context, depositing CMG on dsDNA cannot be used to help the fork overcome DNA damage or to maintain the replisome during fork reversal.

Our nucleotide-resolution analysis of collapsed fork structures identifies at least two DNA processing steps that might be unique to the repair of leading strand collapsed forks (Figure 4D, left). First, the seDSB generated, being almost blunt, must be resected prior to homology-directed strand invasion. Second, on the unbroken sister chromatid, the gap between the 3’ end of the nick and the final primed Okazaki fragment must be filled in to allow strand invasion by the resected end. In contrast, during lag collapse, gap filling is not required, and a ~70 nt ssDNA 3’ overhang is automatically generated (Figure 4D, right) that is probably sufficient to promote strand invasion without further resection (Ira and Haber, 2002; Jakobsen et al., 2019). Our data predict that lead collapse repair requires more processing steps than lag collapse repair and thus might fail in genetic backgrounds that do not support resection (Nacson et al., 2020). In addition, the nearly blunt DNA end generated during lead collapse is an excellent binding site for Ku, suggesting that lead collapse should be more susceptible than lag collapse to NHEJ-dependent chromosomal translocations.

We previously showed that when forks converge during replication termination, CMG is ubiquitylated by CRL2^Lrr1^ and subsequently unloaded by p97. Remarkably, CMG appears to be unloaded by the same mechanism when a single fork encounters a lagging strand nick. This observation argues that CRL2^Lrr1^-dependent CMG unloading operates in multiple cellular contexts and that termination-induced CMG unloading does not require the convergence of two forks. Instead, we recently proposed that CMG ubiquitylation is normally suppressed during elongation by the excluded DNA strand, which prevents CRL2^Lrr1^ binding to the outer face of CMG. In this model, suppression is lost when the excluded strand dissociates from CMG during termination (Low et al., in preparation), or when CMG passes onto dsDNA at a lagging strand nick (Fig 4d). We hypothesize that CMG removal during replication termination and fork collapse serves the same purpose, namely, to prevent the interference of translocating CMGs with transcription and other chromatin-based processes.

## Supporting information

S Video 1 unnicked replication

S Video 2 lead collapse

S Video 3 dCas9 encounter

S Video 4 lag collapse

S Video 5 lag collapse Cul-i

S Video 6 lag collapse p97-i

## Acknowledgement

We thank D. Pellman, J. Haber, and members of the Walter laboratory for helpful discussions and comments on the manuscript. This work was supported by the Howard Hughes Medical Institute and NIH Grants HL098316 and GM80676. K.B.V. and R.A.W. are also supported by an American Cancer Society Postdoctoral Fellowships. G.C. was supported by the Jane Coffin Childs memorial fund. J.C.W. is an investigator of the Howard Hughes Medical Institute.

## Author Contributions

K.B.V. performed all the experiments shown, except for Figure S3b, which was performed by R.A.W. J.M.D. conceived of, developed, and validated the tet-nick system. T.G.W.G. helped validate the tet-nick system. G.C. developed the CMG single molecule imaging approach, and provided labeled GINS, affinity-purified GINS antibody, and Fen1 for the experiments. J.C.W. supervised the work. K.B.V. and J.C.W. wrote the manuscript, with input from the other co-authors.

## Methods

### Preparation of nicked plasmids

The nicked plasmids used for the ensemble experiments were generated from a standard pBlueScript plasmid (pBS), with the following modifications. The pBS BspQI site was removed by site-directed mutagenesis using a QuikChange kit (Agilent). New BspQI sites were added to the plasmids either 245 bp (to visualize the nascent leading strand) or 497 bp (to visualize the nascent lagging strand) away from the intended Tet-nick location using site-directed mutagenesis with the following primer sets: 1) 245 bp, 5’-ATGGTTCACGTAGTGGCTCTTCGCCATCGCCCTGATAGACG-3’ and 5’-GCCACTACGTGAACCATCACCCTAATCAAGTTTTTTGGGG-3’; 2) 494 bp, 5’-CCGAAAAGTGCCACGAAGAGCTGACGCGCCCTGTAGCG-3’ and 5’-CGTGGCACTTTTCGGGGAAATGTGCGCGGAACCCC-3’. A BglII site was added to each plasmid where the Tet-nicks would be inserted using 5’-CCATTCGCCATTCAGAGATCTGCTGCGCAACTGTTG-3’ and 5’-CAACAGTTGCGCAGCAGATCTCTGAATGGCGAATGG-3’ primers. Next the plasmids were linearized with BglII and the first AflII-NheI-TetR-BbvCI-TetR-AvrII-NcoI site was added using Gibson assembly with the following sequence as a duplex (only one strand shown): 5’-CAATTTCCATTCGCCATTCAGACTTAAGGCTAGCTCTCTATCACTGATAGGGACCTCAGCTCTCTATCACTGATAGGGACCTAGGCCATGGAGCTGCGCAACTGTTGGGAAGG-3’. Two additional repeats of these sequences were sequentially added by digesting with AvrII and ligating a NheI-TetR-BbvCI-TetR-AvrII insert into the plasmids. Finally, two 24x LacO arrays were added between the SacI and Kpn sites as previously described (Duxin et al., 2014).

Plasmids were then run on 0.8% agarose gels and the supercoiled DNA was extracted via electroelution. Purified plasmids were nicked with Nb.BbvCI or Nt.BbvCI for leading or lagging strand fork collapse, respectively. The plasmids were then gel purified by electroelution and stored at −20°C in 10 mM Tris, pH 7.5. Plasmid pKV44 has an Nt.BspQI site positioned to nick the nascent leading strand 269 nt away from the first Nb.BbvCI nick site. Plasmid pKV45 has an Nt.BspQI site positioned to nick the nascent lagging strand 515 nt away from the first Nt.BbvCI nick site. pKV44 was used for all ensemble lead collapse experiments except when we wanted to visualize the lagging strand after lead collapse (then we nicked pKV45 with Nb.BspQI). pKV45 was used for all ensemble lag collapse experiments except when we wanted to visualize the leading strand after lag collapse (then we nicked pKV44 with Nt.BspQI). All restriction and nicking enzymes were purchased from New England Biolabs and used according to manufacturer protocols.

### Preparation of egg extracts

The high-speed supernatant (HSS) and nucleoplasmic extracts (NPE) were prepared from *Xenopus laevis* eggs as described previously (Lebofsky et al., 2009).

### Protein expression and purification

Purification of LacR (Dewar et al., 2015) and GINS (Sparks et al., 2019) were described previously. TetR has a His-tag and was purified using Ni-NTA resin as follows. The TetR expression plasmid (*TetR* gene from addgene plasmid 17492 cloned into pET28b) was transformed into BL21 cells in LB supplemented with 50 *μ*g/mL kanamycin. A single colony from this transformation was used to inoculate LB supplemented with 50 *μ*g/mL kanamycin, which was then grown to an OD600 of 0.5 at 37°C. Expression was induced by adding IPTG to 1 mM. After 3 h incubation, the cells were pelleted, and the supernatant was discarded. The cell pellet was resuspended in 2 mL lysis buffer (20 mM Tris, pH 8.0, 1 M NaCl, 5 mM imidazole, 1 mM DTT, 1 cOmplete Protease Inhibitor Cocktail tablet (Roche), 1 mg/mL lysozyme) and rotated for 1 h at 4°C. Lysate was split into two 1.5 mL tubes and centrifuged in a microcentrifuge at 4°C at 13,000 RPM for 30 min. Supernatant was recovered and applied to equilibrated Ni-NTA resin. Samples were spun with Ni-NTA resin for 1 h at 4°C. Resin with lysate was added to a disposable column. Resin was washed twice with 4 mL of wash buffer (20 mM Tris, pH 8.0, 1 M NaCl, 20 mM imidazole, 1 mM DTT). TetR was eluted from column with four, 0.5 mL additions of elution buffer (10 mL, 20 mM Tris, pH 8.0, 1 M NaCl, 1mM DTT, 0.5 M Imidazole). DTT (5 mM) was added to samples immediately after eluting. TetR eluates were combined and dialyzed into 1 L of TetR dialysis buffer (81 mM Tris, pH 7.5, 1.62 mM EDTA, 162 mM NaCl, 1.62 mM DTT) for 2 h at 4°C, then dialyzed into 1 L dialysis buffer overnight. Dialyzed samples were collected and glycerol was added to bring the glycerol to 38% of total volume. Samples were aliquoted and stored at −20°C.

### Ensemble fork collapse reactions

All ensemble replication reactions were carried out as previously described (Lebofsky et al., 2009), with notable changes mentioned below. Briefly, the replication reactions were performed by first pre-binding the plasmid with TetR and LacR, then licensing the DNA in HSS, and finally addition of NPE to initiate replication. One volume of plasmid DNA was pre-incubated with 3 volumes of LacR (23 *μ*M) and 3 volumes of TetR (765 *μ*M) for 20-30 min. After plasmid DNA was pre-bound with LacR and TetR, the DNA was licensed in HSS at a concentration of 6 ng/*μ*L. DNA replication was initiated by mixing 1 volume of licensing mix with 2 volumes of NPE mix that was diluted up to 50% with 1x ELB-sucrose (10 mM HEPES-KOH, pH 7.7, 2.5 mM MgCl_2_, 50 mM KCl, 250 mM sucrose). Reactions were carried out in the presence of [α-P^32^]dATP to radiolabel the replication products. For native agarose gels, reactions were stopped at indicated time points by mixing 1 *μ*L of reaction mix with 6 *μ*L replication stop buffer (8mM EDTA, 0.13% phosphoric acid, 10% ficoll, 5% SDS, 0.2% bromophenol blue, 80 mM Tris-HCl, pH 8.0). Then the proteins were digested with 1 *μ*L of 20 mg/mL Proteinase K for 1 h at 37°C, and plasmids were run on a 0.8-1% agarose gel for ~2.5 h. Gels were then visualized by phosphorimaging on a Typhoon FLA 7000 phosphorimager (GE Healthcare).

### Electron microscopy imaging of DNA structures

Ensemble fork collapse reactions were carried out as described above. After 15 min, 50 *μ*L of reaction mix was stopped by addition to 420 *μ*L extraction stop buffer. The stopped reactions were then RNase and proteinase K treated. DNA was purified by two rounds of phenol-chloroform extraction followed by ethanol precipitation. DNA was resuspended in 20 *μ*L replication stop buffer and was run on a 0.8% agarose gel for 2 h. The gel was then stained with SYBR gold for 1 h to visualize the DNA, and DNA bands of interest were visualized and excised on a blue light box. DNA was electroeluted from the gel slices using an EluTrap system and concentrated with Amicon concentrators (0.5 mL, 100 kDa MWCO).

Five microliters of concentrated DNA were then mixed with 40 *μ*L filtered water and 5 *μ*L 2.5 M ammonium acetate. This mixture was incubated for 10 min. To these mixtures, 2 *μ*L of 0.2 *μ*g/*μ*L cytochrome c solution was added. These solutions were then placed onto Parafilm as drops and allowed to incubate for 15 min. Next, we lightly touched parlodion-coated grids to the surface of the drops, and we dehydrated each grid in 50%, 75%, and 95% ethanol for 15 s each. Each grid was lightly touched to filter paper to dab off excess solution. In order to increase the contrast of the DNA, the grids were rotary shadowed in a Leica ACE600 with a platinum E-beam using the following parameters: 3° angle, 85 W power, 3 × 10^−6^ mbarr, and 3 nm Pt deposition. Finally, we carbon coated the grids to stabilize the parlodion film using the Leica ACE600 carbon E-beam set to the following parameters: 0° angle, 130 W power, 5 ×10^−6^ mbarr, and 2 nm C deposition. Grids were examined with a FEI T12 transmission electron microscope (TEM) equipped with a Gatan 2k SC200 CCD camera at 40 kV or a JEOL 1200EX TEM equipped with a 2k CCD camera (Advanced Microscopy Techniques).

### Nascent strand analysis

To visualize the nascent strands on a denaturing gel, ensemble replication reactions were stopped at indicated time points by mixing 4-6 *μ*L of reaction mix with 30 *μ*L extraction stop buffer (0.5% SDS, 25 mM EDTA, 50 mM Tris-HCl, pH 8.0) followed by addition of 5 *μ*L 4 mg/mL RNase A, and incubation at 37°C for 45 min. Next, the proteins were digested with 5 *μ*L of 20 mg/mL Proteinase K for at least 1.5 h at 37°C. Samples were then diluted to 145 *μ*L with 10 mM Tris, pH 7.5, and DNA was extracted by phenol-chloroform and ethanol precipitation.

Nascent DNA strands were either digested with Nt.BspQI or AflII to visualize the collapse products, as noted in the figure legends. First, ethanol precipitated DNA was resuspended in 5 *μ*L 10 mM Tris, pH 7.5. Each of the digestion reactions was carried out using New England Biolabs enzymes and buffers at a final volume of 10 *μ*L. The Nt.BspQI digestions were carried out with 1 Unit of Nt.BspQI in 1x NEB3.1 Buffer at 50°C for 1 h. The AflII digestions were carried out with 2 Units of AflII in 1x NEB CutSmart Buffer at 37°C for 1.5 h. All reactions were stopped with 5 *μ*L Gel Loading Buffer II (Life Technologies, Cat # AM8547). Digested DNA was incubated at 75°C for 5 min before running on 4% (Nt.BspQI-digested samples) or 10% (AflII-digested samples) polyacrylamide denaturing gels. Gels were dried, exposed to phosphorscreens, and imaged on a Typhoon FLA 7000 phosphorimager (GE Healthcare). The frequency of collapse at each of the collapse sites (Figure 1E) was corrected for the number of adenines expected for each product since products were visualized with [α-32P]dATP.

Sequencing gel ladders were prepared using Thermo Sequenase Cycle Sequencing Kits (USB, Cat # 78500 1KT) with the following primers: 5’-CCATCGCCCTGATAGACGG-3’ (Nt.BspQI, leading strand samples), 5’-TTAAGGCTAGCTCTCTATCACTG-3’ (AflII samples), or 5’-CGAAGAGCTGACGCGCCCTGTAGC-3’ (Nt.BspQI, lagging strand samples). The template DNA was either pKV44 for leading strand analysis or pKV45 for lagging strand analysis.

### Preparation of nCas9 RNP complex

Guide RNA was prepared by annealing AltR CRISPR-Cas9 tracrRNA, ATTO 550 (IDT) with 10-fold excess Alt-R CRISPR-Cas9 crRNA (IDT) in 1x Annealing Buffer (IDT) to yield 4 *μ*M guide RNA (tracrRNA was limiting). The guide RNAs were then frozen at −20°C until needed for experiments. The crRNAs were designed to target specific sequences on the same strand of the 30 kb single molecule DNA (see below). Next, the Cas9 RNP complex was formed by mixing 25 pmol Alt-R S.p. Cas9 H840A Nickase V3 (IDT) with 2 pmol guide RNA in Cas9 binding buffer (20 mM Tris, pH 7.5, 100 mM KCl, 5 mM MgCl_2_, 1 mM DTT, 5% glycerol) and incubating in the dark at room temperature for ~20 min before being used in experiments.

### Single molecule fork collapse reactions

The single molecule replication assay, including flow cell assembly, immunodepletion of GINS from extracts, replication reaction conditions, and image acquisition, was described in detail previously (Sparks et al., 2019). Deviations from the previously published assay are described herein. Coverslips were passivated with 10% Biotin-PEG-SVA and m-PEG-SVA MW5000 (Laysan Bio.). The buffers used to stretch DNA, wash DNA, bind nCas9, and image DNA were degassed for at least 1 h prior to flowing into the flow cell. Flow cells were first incubated with 0.2 mg/mL streptavidin (Sigma) for at least 15 min. Next the flow cells were washed with 500 *μ*L of DNA Blocking buffer (20 mM Tris, pH 7.5, 50 mM NaCl, 2 mM EDTA, 0.2 mg/mL BSA) + 0.5% Tween20 at 500 *μ*L/min. Next, 500 *μ*L of DNA solution containing 67 pg/*μ*L DNA that was biotinylated at each end, DNA Blocking buffer + 0.05% Tween20 and 1.8 mM chloroquine was flowed into the cell at 100 *μ*L/min to double tether the DNA to the coverslip. The flow cell was then equilibrated with 60 *μ*L Cas9 binding buffer (20 mM Tris, pH 7.5, 100 mM KCl, 5 mM MgCl_2_, 1 mM DTT, 5% glycerol) at 20 *μ*L/min. Fifty microliters nCas9 solution was then added at 20 *μ*L/min which contained the 2 nM nCas9 RNP (described above) in the Cas9 binding buffer. Initially, nCas9 bound specifically and nonspecifically across the entire length of the DNA. Site-specifically bound Cas9 molecules remain stably associated with DNA in the presence of 0.5 M NaCl (Sternberg et al., 2014). Therefore, to remove the nonspecifically bound nCas9, the flow cells were washed with 60 *μ*L of Cas9 buffer containing 0.5 M NaCl (20 mM Tris, pH 7.5, 500 mM NaCl, 5 mM MgCl_2_, 1 mM DTT, 5% glycerol) at 20 *μ*L/min. The flow cells were then re-equilibrated with 60 *μ*L Cas9 buffer at 20 *μ*L/min, before adding 30 *μ*L of 200 nM Sytox Green (Thermo Fisher Cat# S7020) in Cas9 binding buffer at 20 *μ*L/min to image the DNA and nCas9. Twenty to fifty fields of view (FOVs) were imaged for both Sytox and nCas9^Atto550^ using alternating 488 nm and 561 nm laser excitation three times (“pre-imaged DNA and nCas9”). Finally, DNA-bound sytox was washed off with 150 *μ*L 1x ELB-sucrose at 10 *μ*L/min. Single molecule replication experiments carried out without nCas9 were prepared using the same procedure except 150 *μ*L of Cas9 binding buffer was flown into the cell at 20 *μ*L/min after the DNA was added, followed by flowing in the Sytox solution, imaging the DNA, and flowing in the 1x ELB.

Endogenous GINS was immunodepleted in two rounds from HSS and three rounds from NPE (1 h each) at 4°C. The depleted extracts were used to make licensing, initiation, and replication mixes as previously described (Sparks et al., 2019) at room temperature. The double-tethered, nCas9-bound DNA was licensed by flowing in 20 *μ*L of GINS-depleted HSS licensing mix at 10 *μ*L/min and incubating for 4-15 min. Replication was then initiated with 20 *μ*L of GINS-depleted HSS/NPE initiation mix that included 0.01 mg/mL recombinant GINS^AF647^, 2 *μ*M Fen1-mKikGR D179A, and 3.7 nM nCas9^Atto550^ at 10 *μ*L/min. After 4 min, 50 *μ*L of a new GINS-depleted HSS/NPE replication mix was flown in at 10 *μ*L/min that included 2 *μ*M Fen1-mKikGR D179A, but did not include GINS^AF647^ or nCas9^Atto550^ to remove fluorescence background from excess proteins. The absence of nCas9 in the final replication mix prevents rebinding of nCas9 during the collapse reactions. Where indicated, p97-i (NMS-873, Sigma) or Cul-i (MLN4924, Active Biochem) inhibitors were added to the initiation and replication mixes at a final concentration of 200 *μ*M. Images of Fen1-mKikGR, nCas9^Atto550^, and GINS^AF647^ were acquired every minute for 1 h by cycling among the 488 nm (64-65° TIRF angle, 0.23 mW, 100 ms exposure, 999 EM GAIN), 561 nm (61-63° TIRF angle, 0.35 mW, 100 ms exposure, 999 EM GAIN), and 647 nm (61-63° TIRF angle, 0.15 mW, 100 ms exposure, 999 EM GAIN) lasers at each of the fields of view. Specific microscope configurations were previously described (Sparks et al., 2019). Movies were collected using NIS Elements software and saved as nd2 files.

### Single molecule data analysis

All image analysis was performed using a combination of freely available and custom Matlab scripts. Movie files were imported into a graphical user interface (GUI) using Bio-Formats to convert the nd2 files into readable metadata and matrices (Linkert et al., 2010). The movies were then stabilized to remove drift using a Matlab script for efficient subpixel image registration (Guizar-Sicairos et al., 2008). The nCas9 was targeted to the DNA asymmetrically to divide the DNA into short and long arms relative to the nicking site. To determine which side of the DNA the nCas9 was binding to, the nCas9 channel from the “pre-imaged DNA and nCas9” (described above) with the nCas9 channel from the replication movie was aligned using the same subpixel image registration script. The brightness and contrast of the movies were adjusted for each channel in Matlab. The max projection of the CMG signal was overlaid with the image of the unreplicated DNA (acquired before addition of extract) to identify which DNA molecules were replicated. Then DNA molecules of interest were manually selected to assemble kymographs. Molecules were chosen based on three criteria: 1) multiple DNA molecules did not overlap significantly with each other, 2) nCas9^Atto550^ signal was correctly positioned on the DNA, and 3) the DNA appeared to be nearly fully stretched. The kymographs were then categorized according to the type of events observed, as shown in Figures S2F, S2G, and S2H. CMG velocities were determined from the linear fit of the center position of the GINS^AF647^ signal over time. These signal positions were determined using a modified version of the u-track Matlab software (Jaqaman et al., 2008). The time to unload CMG after collapse was determine by counting the number of frames until GINS^AF647^ signal was lost (1 min each) after nCas9^Atto550^ signal was lost. The time to unload CMG was counted as zero minutes for the events in which CMG and nCas9 appeared to be lost at the same time.

### Quantification and statistical analysis

All ensemble experiments were repeated three or more times except for Extended Data 3b, which was done twice. All single molecule experiments were performed two or more times. All image analysis, including replication and sequencing gels and single molecule movies, was performed using Matlab. The Kaplan-Meier curve in Figure S2E was generated using Matlab’s built-in empirical cumulative distribution function (ecdf). Error bars were generated from three individual biological replicates. P values were calculated using Matlab’s two-sample *t*-test function. Additional relevant statistical details are mentioned in the figure legends.

### Code and data availability

All custom-written Matlab code and raw data will be made available upon request.

---

**Figure S1.**
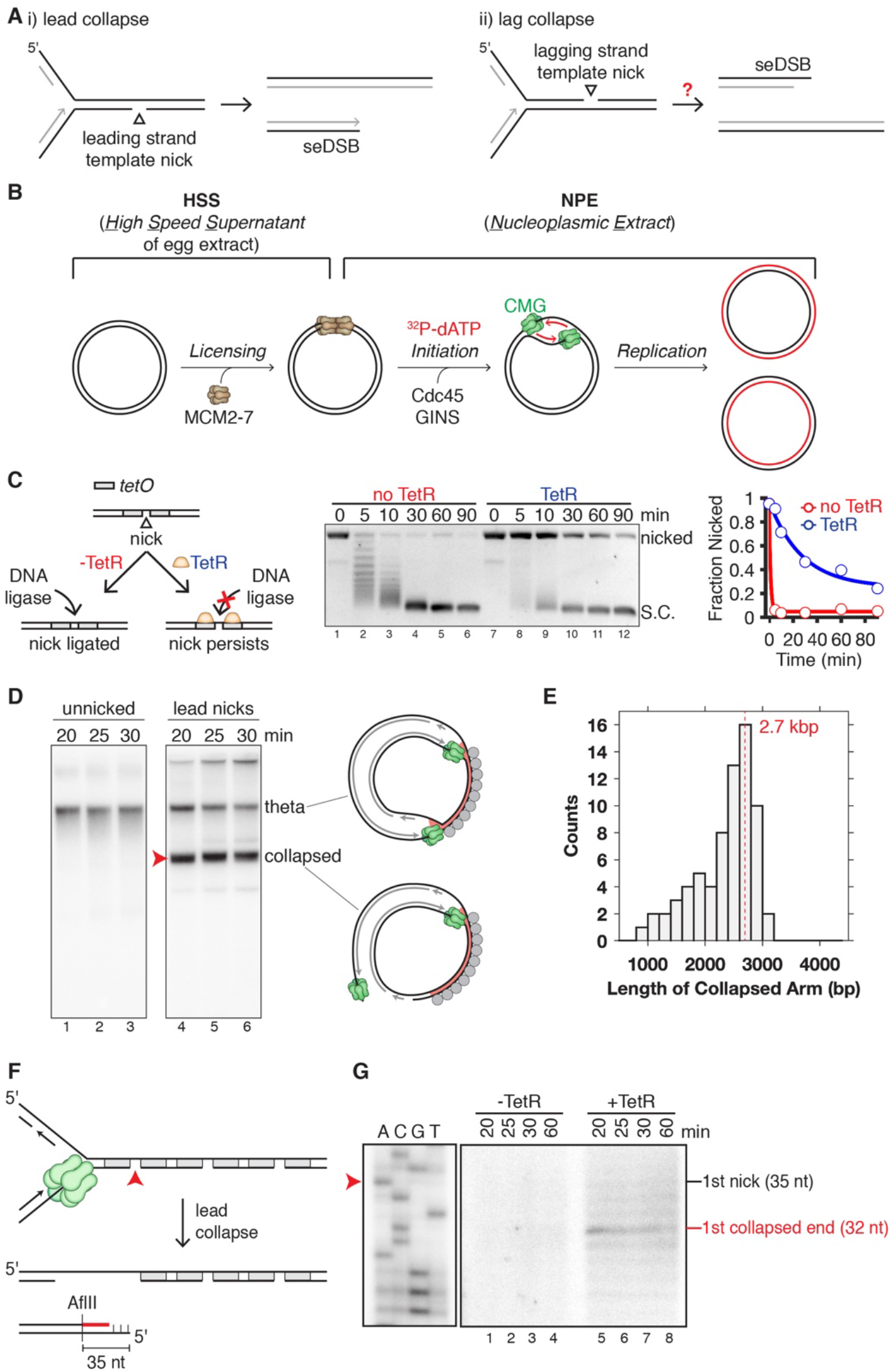
Leading strand fork collapse in *Xenopus* extracts using TetR-nick protection strategy. (A) General representations of lead and lag fork collapse. (B) Experimental workflow for DNA replication in *Xenopus* egg extracts. DNA is first licensed in HSS to load the MCM2-7 double hexamer, followed by addition of NPE to initiate a single round of DNA replication. (C) TetR-nick protection strategy. In the absence of TetR, nicks are ligated by DNA ligase (left branch of cartoon, lanes 1-6, and red line in graph). In the presence of TetR, nicks are much more stable, presumably because TetR prevents access of DNA ligases to the nicks (right branch of cartoon, lanes 7-12, and blue line in graph). (D) Unnicked plasmid or plasmid with a bottom strand nick (Lead collapse) was replicated in the presence of TetR and LacR using egg extracts (depicted in Figure 1A). (E) Length distribution of the collapsed arms visualized by EM. The theoretical length of the collapsed arm after lead collapse is 2.7 kbp. Length is estimated assuming B-form DNA with a rise per basepair of 0.34 nm. (F) Schematic of the nascent leading strand products generated from lead collapse at the first nick. The red line depicts the newly synthesized leading strand after digestion by AflII, which cuts 35 nt from the nick. (G) Plasmid with bottom strand nick (lead collapse) was replicated in egg extract in the presence and absence of TetR, as indicated. Nascent leading strand products were analyzed on a urea PAGE gel after lead collapse and AflII digestion. In the absence of TetR, nicks are ligated, explaining the lack of collapse products. The prominent leading strand product (32 nt) is three nt shorter than the terminal AflII fragment (35 nt).

**Figure S2.**
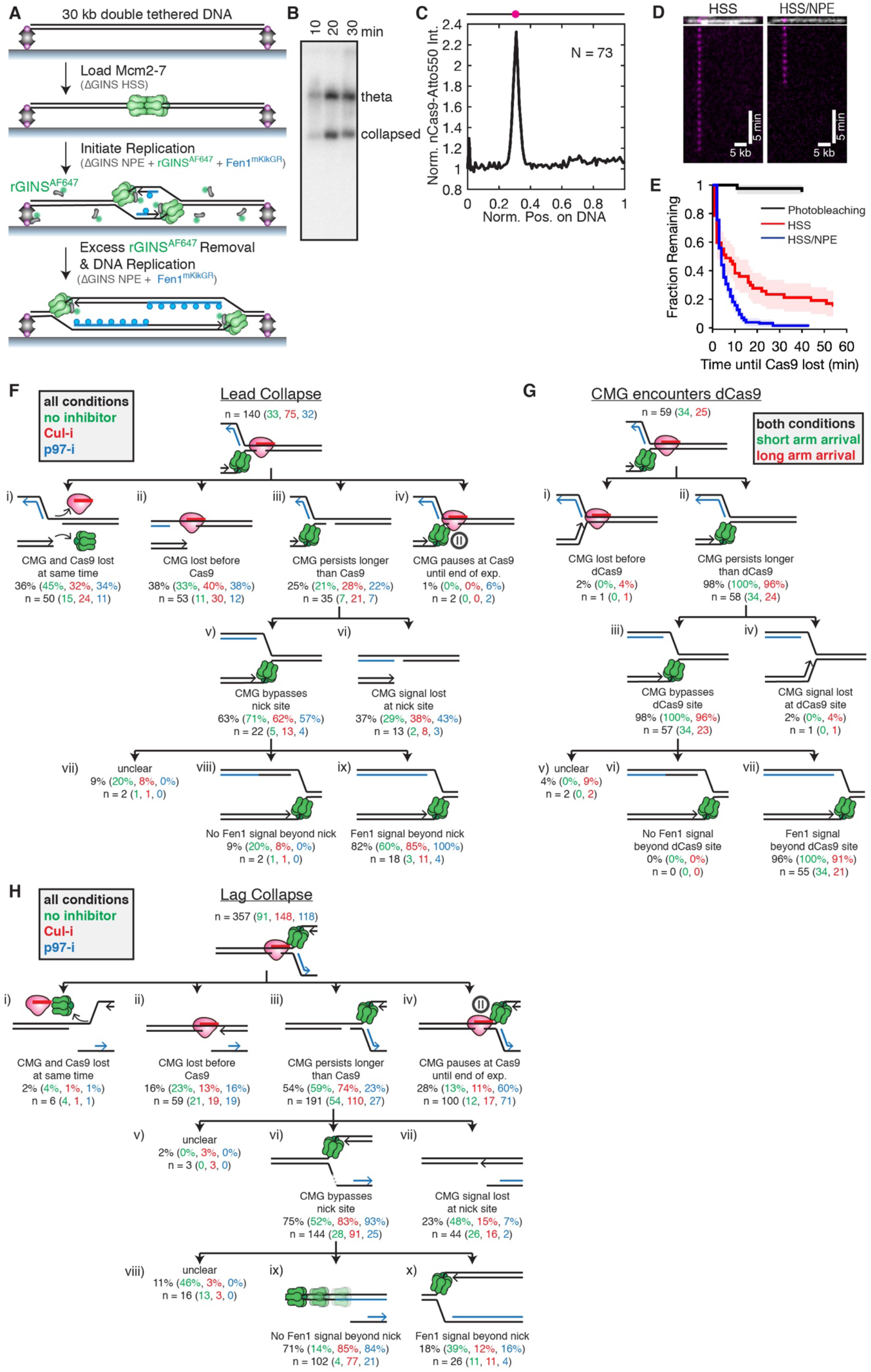
Single molecule visualization of CMG and nCas9 on double-tethered DNA. (A) Schematic of single molecule replication assay. First, double-biotinylated 30 kb DNA is stretched and double-tethered to a streptavidin-coated coverslip. Next, GINS-depleted HSS is injected into the flow cell to load MCM2-7 double hexamer (licensing). Replication is then initiated with GINS-depleted NPE supplemented with recombinant GINS tagged with Alexa Fluor 647 (rGINS^AF647^, gray shape with green circle) and Fen1^mKikGR^ (blue circle). After 4 minutes, excess GINS is removed by flowing in GINS-depleted NPE supplemented only with Fen1^mKikGR^. Fen1^mKikGR^ delays the dissociation of PCNA from DNA and therefore appears to bind PCNA molecules that are deposited during Okazaki fragment synthesis, leading to labeling of the entire replication bubble. (B) nCas9-induced replication fork collapse. The experiment is similar to the one shown in Figure S1D, except that nCas9 H840A is used to nick the DNA. (C) Position of nCas9^Atto550^ signal on double-tethered DNA. The black line above the plot shows the theoretical position for nCas9 binding (magenta dot). Only a single nCas9 was targeted to the DNA for this experiment. (D) Representative kymographs of nCas9 binding to DNA in HSS or 1:2 mixtures of HSS and NPE which mimic the condition used during single molecule DNA replication. The DNA (white) and nCas9 (magenta) shown above each kymograph are from imaging the nCas9 and DNA before adding the extracts. (E) The rates of nCas9 loss in HSS or HSS/NPE compared to the photobleaching rate. Shaded regions represent 90% confidence bounds. (F, G, and H) Breakdown of all categories of events observed during (F) lead collapse, (G) CMG collision with dCas9, or (H) lag collapse in single molecule assays. Below each category, the number of events observed in that category under three different conditions is listed in brackets. For F and H, no drug, green; Cul-i, red; p97-i, blue. For G, CMG collides with dCas9 from the short arm, green; CMG collides with dCas9 from the long arm, red. Black, sum of all conditions in a category. Cartoons only depict arrival from the short arm. Percentages are calculated for each tier of the hierarchy.

**Figure S3.**
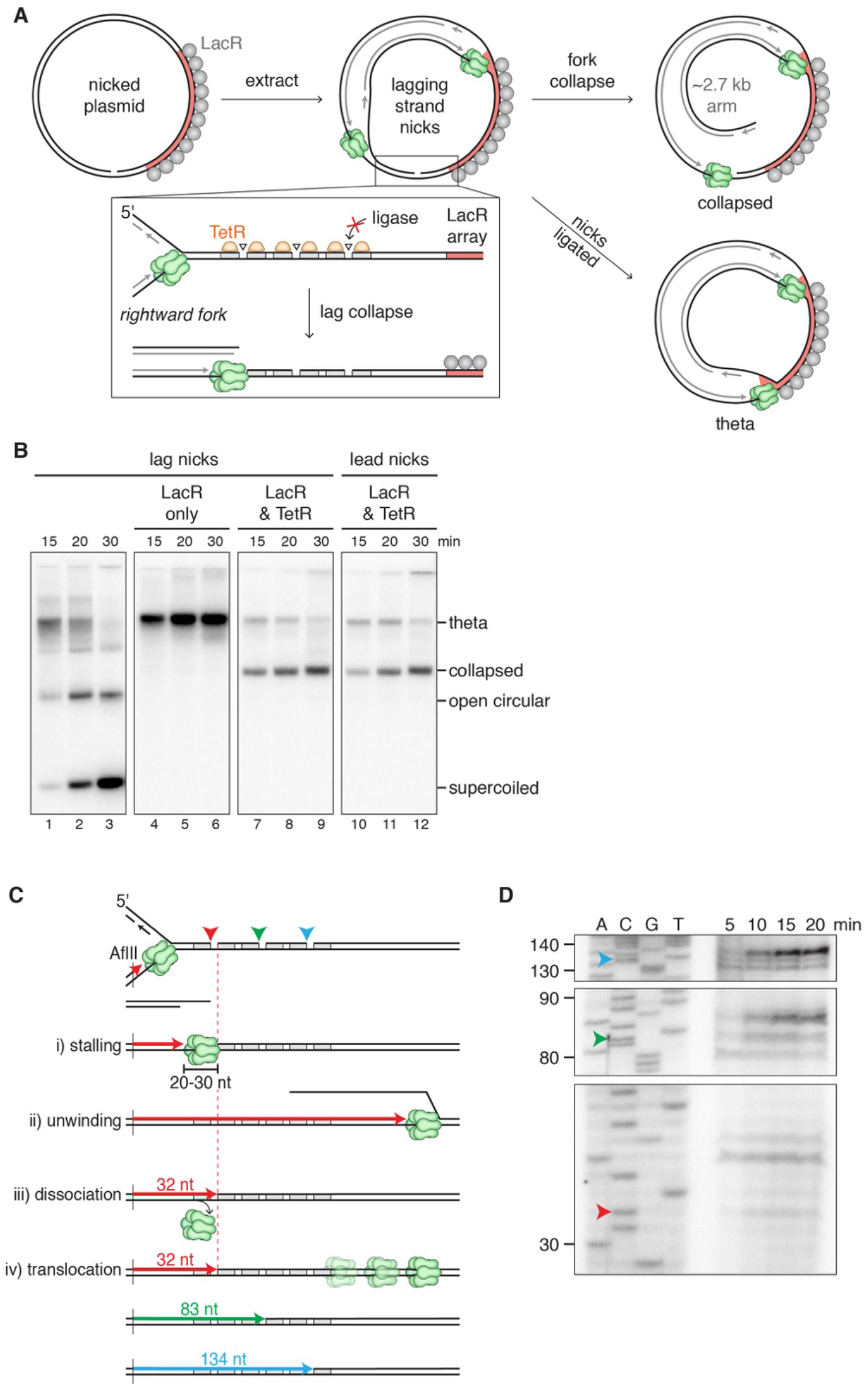
Lagging strand fork collapse in *Xenopus* extracts using the Tet-nick strategy. (A) Depiction of structures generated during replication of a top strand (lag collapse) plasmid in the presence of TetR. Inset, detailed representation of the three nicks and flanking *tetO* sites. (B) Plasmid with a bottom strand (lead) nick or top strand (lag) nick was replicated in the presence of LacR +/−TetR using egg extracts. In the absence of TetR and LacR (lanes 1-3), nicks were ligated, and replication went to completion, generating the expected open circular and supercoiled products. In the presence of LacR, but no TetR (lanes 4-6), the unprotected nicks were ligated, and a prominent theta band was generated from forks converging on the LacR array (as in (A), bottom route). In the presence of both LacR and TetR (lanes 7-9), the nicks were protected, and lag collapse occurred (as in (A), top route), generating the same collapsed product detected from lead collapse (lanes 10-12 and Figure S1D). (C) Same as Figure 3B, except that lead products for all three nicks are shown for model iv. (D) Same as Figure 3D, except that DNA was digested with AflII (35 nt away from first nick) and run on a 10% polyacrylamide gel to improve the resolution. A significant fraction of the leading strands were extended approximately 3 nt beyond the nick site, consistent with limited strand displacement synthesis.

**Figure S4.**
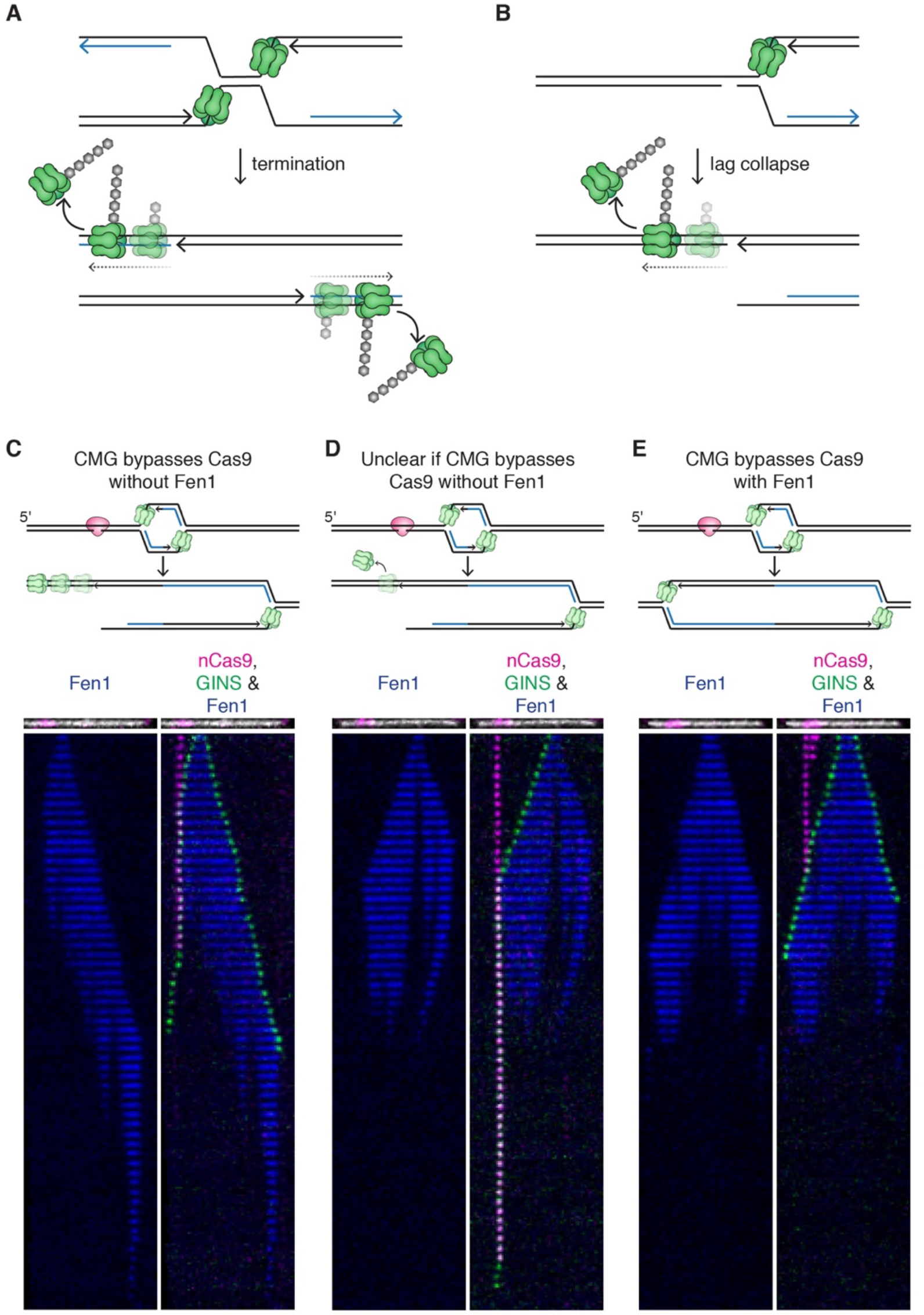
Parallels between termination and collapse, and representative kymographs of different classes of “lag collapse” events. (A) During replication termination, replication forks converge, CMGs pass each other and pass onto dsDNA when they hit the nascent lagging strand of the opposing fork, followed by CRL2^LRR1^-dependent CMG ubiquitylation and extraction by the p97 ATPase (Dewar et al., 2017). (B) During lag collapse, CMG passes onto dsDNA when it hits the nick in the lagging strand template, followed by cullin-RING and p97-dependent chromatin extraction. (C) Representative single molecule kymograph and interpretive cartoon of lag collapse events in which CMG translocated unidirectionally away from the collapse site after collapse (quantified in Figure S2H,ix). The DNA (white) and nCas9 (magenta) shown above each kymograph are from imaging the nCas9 and DNA before extract addition. (D) Representative kymograph of lag collapse events in which CMG was unloaded too rapidly to assess whether Fen1 signal trailed behind CMG (quantified in Figure S2H,viii). (E) Representative kymograph of events in which CMG bypassed nCas9 with a trailing Fen1 signal. This signature suggests that the fork did not collapse, presumably because nCas9 did not nick the DNA or because the nick was ligated prior to CMG encounter (quantified in Figure S2H,x).

## Supplemental Information

**Supplemental Video 1. Replication of unnicked DNA.** Replication of a single, double tethered DNA was visualized by TIRF microscopy. CMG shown in green, Fen1^mKikGR^ in blue. Frames collected every minute for 1 h. Corresponding kymograph in Figure 2A,i.

**Supplemental Video 2. Leading strand fork collapse.** Collision of a replication fork arriving from the short arm with nCas9 results in leading strand fork collapse. CMG shown in green, Fen1^mKikGR^ in blue, nCas9 in magenta. Frames collected every minute for 1 h. Corresponding kymograph in Figure 2A,ii.

**Supplemental Video 3. Replication collision with dCas9.** Unlike leading strand fork collapse, in which CMG is lost at the nCas9, CMG bypasses dCas9 and replication continues. CMG shown in green, Fen1^mKikGR^ in blue, dCas9 in magenta. Frames collected every minute for 1 h. Corresponding kymograph in Figure 2A,iii.

**Supplemental Video 4. Lagging strand fork collapse.** Collision of a replication fork arriving from the long arm with nCas9 results in lagging strand fork collapse. CMG shown in green, Fen1^mKikGR^ in blue, dCas9 in magenta. Frames collected every minute for 1 h. Corresponding kymograph in Figure 4A,Veh.

**Supplemental Video 5. Lag collapse in the presence of Cul-i.** CMG shown in green, Fen1^mKikGR^ in blue, nCas9 in magenta. Frames collected every minute for 1 h. Corresponding kymograph in Figure 4A,Cul-i.

**Supplemental Video 6. Lag collapse in the presence of p97-i.** CMG shown in green, Fen1^mKikGR^ in blue, nCas9 in magenta. Frames collected every minute for 1 h. Corresponding kymograph in Figure 4A,p97-i.

